# Variation in genome architecture and epigenetic modification across the microsporidia phylogeny

**DOI:** 10.1101/2025.01.20.633886

**Authors:** Pascal Angst, Dieter Ebert, Peter D. Fields

**Author notes:** Corresponding author Pascal Angst Department of Environmental Sciences, Zoology University of Basel Vesalgasse 1, 4051 Basel.

## Abstract

Microsporidia are a model clade for studying intracellular parasitism, being well-known for their streamlined genomes and their extreme life history. Although microsporidia are highly diverse and ecologically important to a broad range of hosts, previous research on genome architecture has focused primarily on the mammal-infecting genus *Encephalitozoon*. Here, we expand that work, testing the universality of the patterns observed in *Encephalitozoon* by investigating and comparing variation in genetic and epigenetic architectures in the high-quality genome assemblies of several major microsporidia clades. Our comparison of nine genomes, including the first genome assemblies of *Binucleata daphniae*, *Gurleya vavrai*, and *Conglomerata obtusa*, and revised, improved assemblies of *Glugoides intestinalis* and *Ordospora colligata*, found limited conservation of genetic and epigenetic architecture across all microsporidia, although many genomic characteristics, such as nucleotide composition and repeat content, were shared between genomes of the same or related clades. For example, rRNA genes were hypermethylated in all species, but their position close to chromosome ends was only found in the *Encephalitozoon* and its sister clade. GC-content varied widely, linked to genome size, phylogenetic position and activity of repeat elements. These findings enhance our insight into genome evolution and, consistent with findings from other systems, suggest epigenetic modification as a regulatory mechanism of gene expression and repeat element activity in microsporidia. Our comparative genome analysis reveals higher variation among microsporidia than previously supposed.

## Introduction

The study of parasite evolution looks at the mechanisms by which parasites adapt to exploit their hosts (Poulin & Randhawa, 2015; Schmid-Hempel, 2021). These adaptations include physical structures used to invade or attach to the host, secretory molecules that subvert the host’s immune system or alter its behavior to the parasite’s advantage, and genomic rearrangements that optimize gene transcription and genome replication. Genome evolution particularly has received much recent attention in the field of microsporidia. These fungi-related intracellular parasites exhibit high variation in genome length, possibly related to their life history (Haag et al., 2020), but although comparative studies of microsporidia have provided initial insights into what is arguably the most extreme form of parasitism, including the molecular basis of parasitic adaptations (Nakjang et al., 2013; Wadi & Reinke, 2020; Williams et al., 2022), the lack of high-quality genomes for most major microsporidia clades in this little understood branch of the tree of life has limited comparative studies of genome architecture to single genera (see e.g., Khalaf et al. (2024), Mascarenhas dos Santos et al. (2023)). The available short-read-based genome assemblies have provided inadequate insights into microsporidia genomic architecture, despite hinting at its high diversity.

Microsporidia exhibit highly derived genomic features for parasitism, including mitochondrion genome loss and nuclear genome compaction (Jespersen et al., 2022), the latter of which is highly variable among microsporidia clades, with different forms and levels of (non-)coding DNA loss. Extreme examples can be seen in the mammalian parasite genus *Enterocytozoon*, which lacks most genes involved in glycolysis and is therefore energetically completely dependent on its host (Wiredu Boakye et al., 2017), and in the Encephalitozoonidae family, which has reduced inter- and intragenic regions, including repeat content (Corradi & Slamovits, 2011; Mascarenhas dos Santos et al., 2023). An opposite phenomenon, genome expansion, is observed secondarily in microsporidia with mixed-mode (horizontal and vertical) transmission, such as *Nosema bombycis* and *Hamiltosporidium tvaerminnensis* (de Albuquerque et al., 2020; Parisot et al., 2014). This likely occurs because vertical transmission reduces effective population size and thus the efficacy of natural selection to limit genome expansion, e.g., through the proliferation of transposable elements (Haag et al., 2020). While the quality of microsporidian genome assemblies is not as essential when it comes to comparing the length or presence of genetic elements, we do need high-quality assemblies to study the abundance and relative location of these genetic elements across species.

Although microsporidia are found in a wide range of host species, including humans and agriculturally important animals, from different environments, most microsporidia seem to occur in aquatic environments with the greatest richness found in Crustacea (Bojko & Stentiford, 2022). Indeed, thanks, perhaps, to the vast abundance of Crustacea and the opportunistic nature of microsporidia (Weiss & Becnel, 2014), Crustacea host microsporidia from all major clades (Bojko et al., 2022). Additionally, because many microcrustaceans such as the model system *Daphnia* are filter feeders, they accumulate infectious pathogen cells during feeding, which provides the initial contact point and opportunity for microsporidia–host interactions. *Daphnia* alone are known to be regularly infected by species from at least four of the seven major microsporidia clades, as well as an expanded microsporidium that still has a mitochondrion (Ebert, 2008; Haag et al., 2014).

In this comparative genomic study, we aim to better understand genomic evolution in microsporidia by elucidating the causes of variation in their genomic features, and the arrangement and epigenetic modifications of their genome. Only recently, and only for a few microsporidia genera, have genomic architecture and methylation patterns been described. For example, a recently-described genomic architecture for three species in the human-infecting genus *Encephalitozoon* found that each chromosome in the genomes of *Encephalitozoon* spp., from the ends to the core, consists of 5-mer telomeric repeats, telomere-associated repeat elements (TAREs), hypermethylated ribosomal RNA (rRNA) genes, less methylated subtelomeres, and a hypomethylated chromosome core (Mascarenhas dos Santos et al., 2023). Our aim is to evaluate the generality of this architecture by using high-quality genome resources to examine microsporidia from diverse clades of the phylum. We look at *Encephalitozoon intestinalis*, its close relative *Ordospora colligata*, *Vairimorpha necatrix* also of the Nosematida clade, *Glugoides intestinalis* of the sister clade Enterocytozoonida, *Hamiltosporidium tvaerminnensis* of the orphan clade, three microsporidia of the Amblyosporida clade (*Binucleata daphniae*, *Gurleya vavrai*, and *Conglomerata obtusa* (≡ *Larssonia obtusa*)), and one expanded microsporidium, *Mitosporidium daphniae*. We further explore the processes that drive genome rearrangements and epigenetic modifications in these microsporidia. Previous studies have shown a relationship between mode of transmission and repeat content in the microsporidia genome (de Albuquerque et al., 2020; Haag et al., 2020), suggesting that, for example, life history contributes to the evolution of genomic architecture. As part of our study on genome organization and (epi)genetic features in these diverse microsporidia, we also contribute here the first high-quality genome assemblies and epigenetic annotations for a number of understudied microsporidia clades.

## Results

### Genome assemblies

We generated genome assemblies for five microsporidia (Table 1), including first-time assemblies for *B. daphniae*, *G. vavrai*, and *C. obtusa*, as well as enhanced assemblies for *O. colligata* and *G. intestinalis*. To expand our taxon sampling, we also included available high-quality genomes of four other microsporidia: *E. intestinalis* (Mascarenhas dos Santos et al., 2023), *H. tvaerminnensis* (Angst et al., 2023), *M. daphniae* (Angst et al., in prep.), and *V. necatrix* (Svedberg et al., 2024) (Table 1). The overall length of the nine genome assemblies correlated negatively with their GC-content (*F*(1,7) = 19.44, *P* = 0.003, *R*^2^ = 0.74; Figure 1), following the trend seen in all (draft) microsporidia reference genome assemblies available in the NCBI database (Figure S1, Table S1). The Amblyosporida, *H. tvaerminnensis*, and *V. necatrix* genomes had about 15 % lower GC-content (22 – 29 %) and were about three times longer (10 – 22 Mb) than the smaller *E. intestinalis*, *O. colligata*, *G. intestinalis*, and *M. daphniae* genomes (34 – 43 % and 3 – 6 Mb). The larger genomes had a higher repeat content (26 – 50 %) and a higher number of protein-coding genes (2,847 – 4,231 bp) than the smaller genomes (12 – 20 % and 1,768 – 2,840 bp), except for *G. intestinalis*, whose genome consisted of 43 % repeats (Positive correlation between assembly and repeat length: *F*(1,7) = 36.35, *P* = 0.001, *R*^2^ = 0.84). The protein-coding genes in larger genomes tended to be shorter on average (952 – 1,081 bp) than in shorter genomes (1,019 – 1,447 bp), except for *H. tvaerminnensis* (1,391 bp) whose protein-coding genes were closest to the expanded microsporidium *M. daphniae* (1,447 bp) in average length.

**Figure 1:**
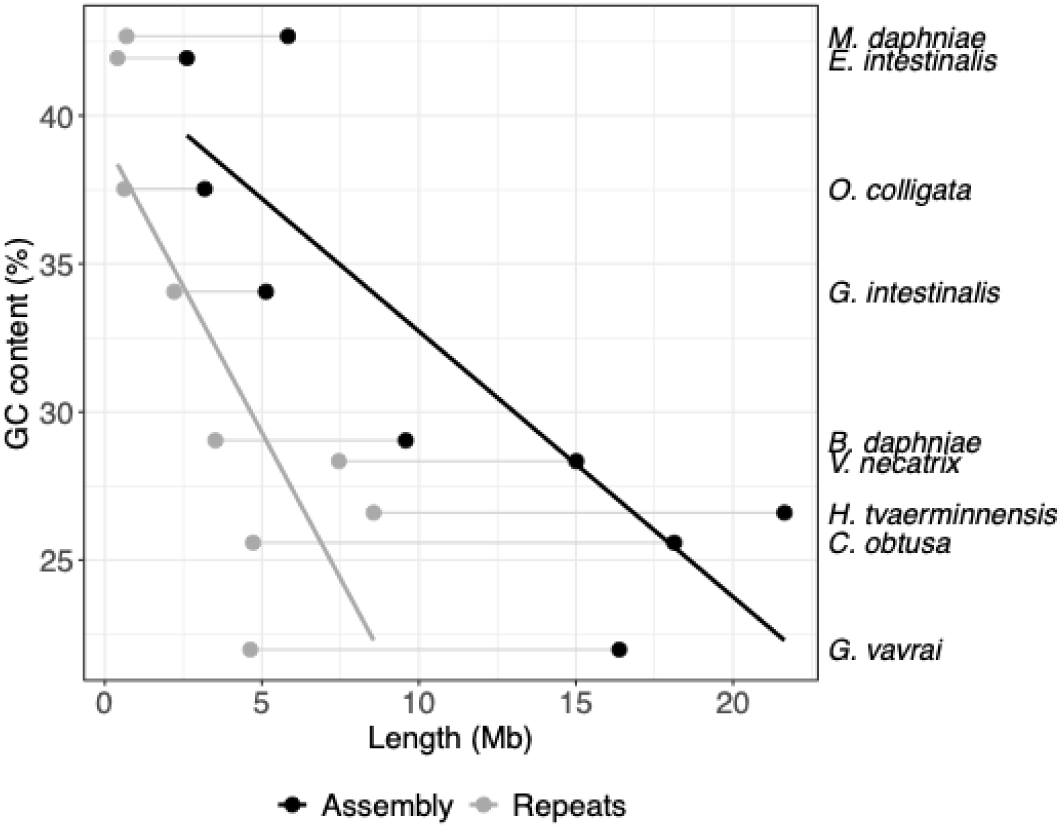
Relationship between genome length, total repeat length, and GC-content in nine microsporidia species. Assembly length (black dots and line) and total length of repeats (gray dots and line) correlated negatively with GC-content. Species names are displayed to the right of the plot at the corresponding GC-content.

**Table 1:** Assembly information for the analyzed m 151 icrosporidia genomes. BUSCO completeness was calculated using the microsporidia specific microsporidia_odb10 database, which is not specific to expanded microsporidia like *M. daphniae*.

| *Glugoides intestinalis* | *Gurleya vavrai* | *Hamiltosporidium tvaerminnensis* | *Mitosporidium daphniae* | *Ordospora colligata* | *Vairimorpha necatrix* |
| --- | --- | --- | --- | --- | --- |
| 5.13 | 16.37 | 21.64 | 5.83 | 3.18 | 15.01 |
| 34.06 | 21.97 | 26.6 | 42.68 | 37.53 | 28.34 |
| 439,321 | 989,478 | 1,444,059 | 864,371 | 374,328 | 1,281,651 |
| 13 | 20 | 17 | 10 | 9 | 12 |
| 83.8 | 90.3 | 94 | 37.6 | 99 | 96.8 |
| 2.21<br>43.06 | 4.63<br>28.27 | 8.56<br>39.56 | 0.69<br>11.79 | 0.62<br>19.54 | 7.45<br>49.68 |
| 2,331 | 3,537 | 3,573 | 2,840 | 1,768 | 2,971 |
| 1,127 | 978 | 1,391 | 1,447 | 1,114 | 1,081 |
| PacBio HiFi | ONT plus Illumina | PacBio CLR plus ONT | ONT plus Illumina | PacBio HiFi | PacBio HiFi |
| new | new | Angst et al. (2023) | Angst et al. (in prep.) | new | Svedberg et al. (2024) |

|  | *Binucleata*<br>*daphniae* | *Conglomerata*<br>*obtusa* | *Encephalitozoon*<br>*intestinalis* |
| --- | --- | --- | --- |
| Total length (Mb) | 9.58 | 18.13 | 2.61 |
| GC-content (%) | 29.04 | 25.59 | 41.94 |
| Contig N50 (bp) | 332,081 | 6,631 | 230,490 |
| Contig number | 31 | 5,653 | 11 |
| BUSCO completeness (%) | 85.7 | 88.5 | 99.5 |
| Total length of repeats (Mb and %) | 3.52<br>36.79 | 4.72<br>26.02 | 0.4<br>15.18 |
| Number of protein-coding genes | 2,847 | 4,231 | 2,074 |
| Mean len. of protein- coding genes (bp) | 1,049 | 952 | 1,019 |
| Sequencing technology | ONT plus Illumina | Illumina | ONT plus Illumina |
| Publication | new | new | Mascarenhas dos Santos et al. (2023) |

Our improved *O. colligata* and *G. intestinalis* genome assemblies were more complete than previous assemblies, containing subtelomeric sequences for most chromosomes (Table S2). Phylogenetic inference confirmed previous rRNA gene phylogenies, placing our eight core microsporidia genomes into four different clades; *M. daphniae* was placed with the expanded microsporidium *Paramicrosporidium saccamoebae* (Figure 2 and S2). Our first high-quality genome assemblies in the Amblyosporida clade were all from the Gurleyidae family.

**Figure 2:**
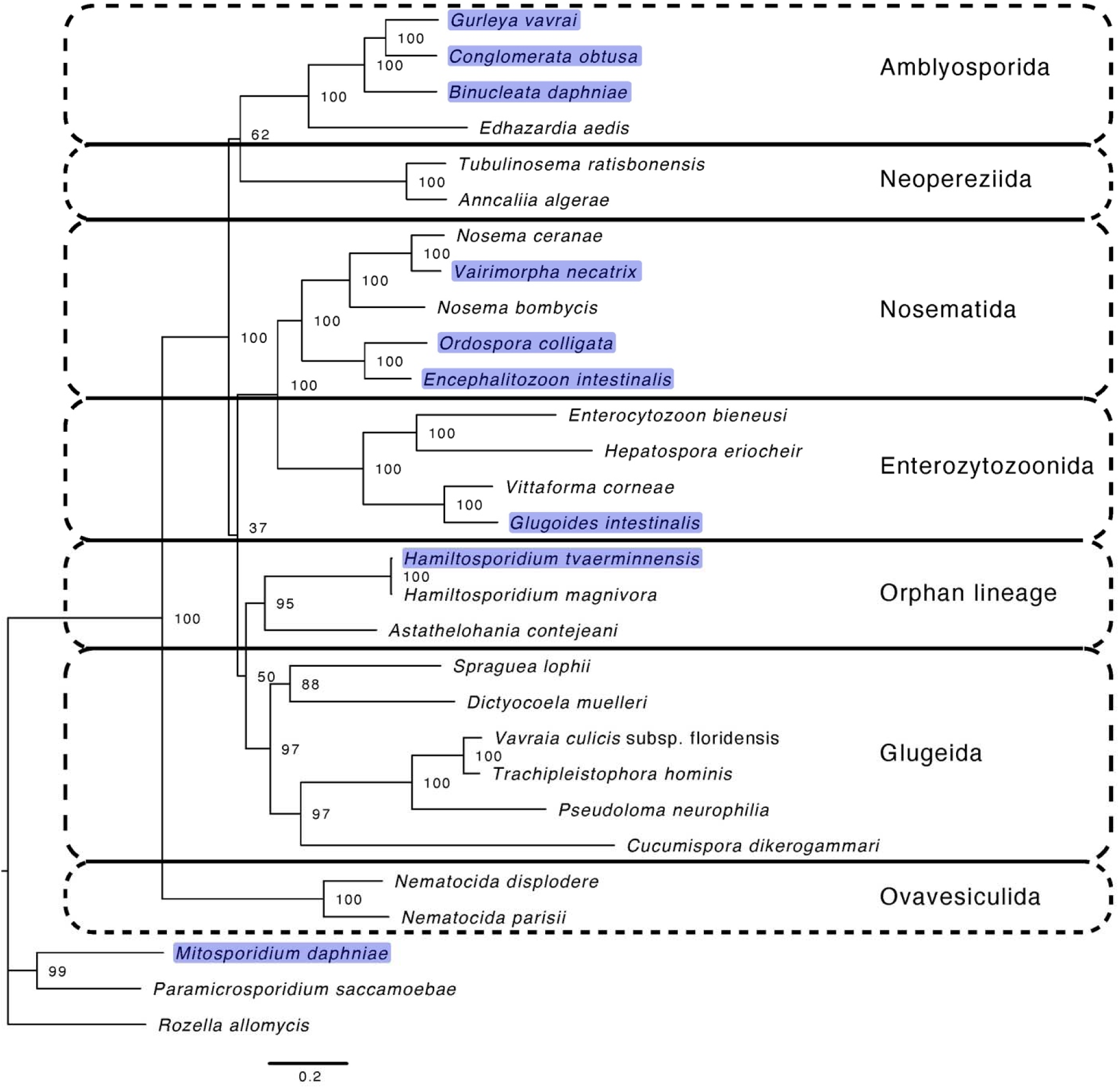
Maximum likelihood phylogeny of the core and expanded microsporidia with *Rozella allomycis* as an outgroup. In addition to the species discussed here (highlighted), we selected a representative sample of microsporidia species with available data for phylogenetic inference based on 15 single-copy orthologs. Node labels represent bootstrap values from 1,000 runs, and major microsporidia clades are delineated according to Bojko et al. (2022).

### Genome architecture and synteny

Our reference and starting point for this comparison of microsporidia genome architecture was the well-described chromosome arrangement of *Encephalitozoon* (Mascarenhas dos Santos et al., 2023). We first compared *E. intestinalis* to its close relative *O. colligata*, finding them to have a similar genome architecture with a 5-mer telomeric repeat (TTAGG) followed by a large and a small rRNA gene each (= rRNA operon), which we found on both ends of six out of the nine contigs and on one end of each of the other three contigs (Figure 3, Table S2). The next most closely related species, *V. necatrix* and *G. intestinalis*, had the same 5-mer telomeric repeat and a 4-mer telomeric repeat motif (TTAG), respectively. In *G. intestinalis*, rRNA genes followed the telomeric repeats on both ends of four contigs, on one end of seven contigs, and were absent in two contigs with telomeric repeats, indicating variation in chromosomal arrangement. The *V. necatrix* genome was more repetitive than its closest relatives, with more repeats between telomeres and rRNA genes, and with variation in their presence and number. In the other major clades containing Amblyosporida and *H. tvaerminnensis*, the genome architecture differed even more widely. The rRNA operons were not bound to the subtelomeres and occurred in tandem with tail-to-tail orientation. *Hamiltosporidium tvaerminnensis* had ten rRNA operons at five locations, and *G. vavrai* had 14 copies at seven locations. Each pair of juxtaposed rRNA operons surrounded specific LINE repeats that were predominantly found there. *G. vavrai* was also found to contain the previously described telomeric repeat motif of *H. tvaerminnensis* (TTAGGG; Angst et al. (2023)). From the less contiguous genome assemblies of *B. daphniae* and *C. obtusa*, we could recover no telomeric repeats and only a few to no rRNA genes.

**Figure 3:**
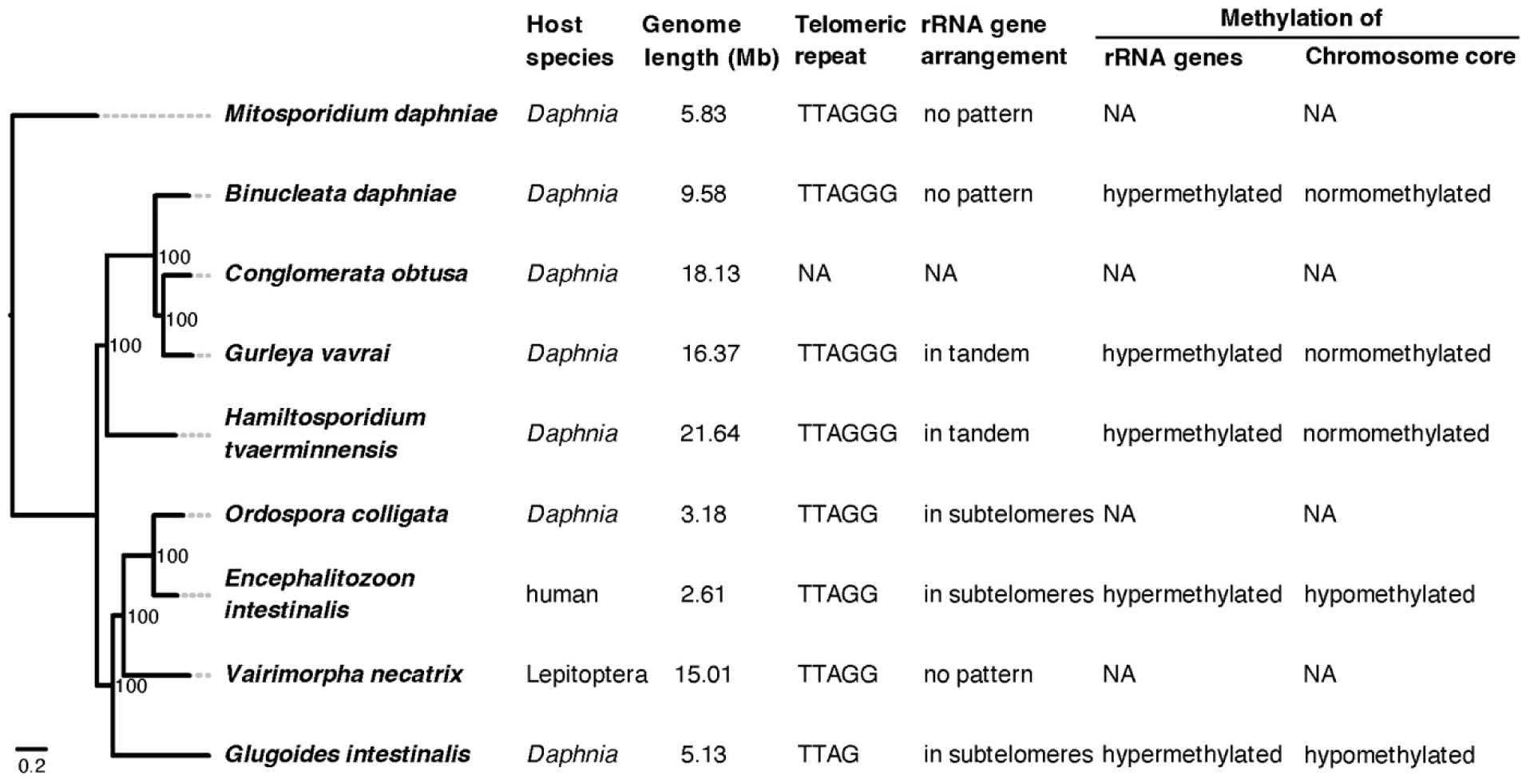
Summary of (epi)genetic features across species. Each row in the maximum likelihood phylogeny corresponds to a species and its associated (epi)genetic features. Node labels represent bootstrap values from 1,000 runs.

Overall, we found limited concordance in sequence arrangement (or synteny) across the analyzed microsporidia unless they were closely related (Figure 4). For example, although single-copy BUSCO genes, which are preserved due to their importance for organismal function and survival, occurred in conserved linkages across closely related species, only a few BUSCO linkage groups were shared by all core microsporidia. This was as expected due to the high degree of species divergence. Throughout the phylogeny, we found no centromeric regions based on chromosome-wide GC or repeat content, supporting hypotheses that microsporidian chromosomes might be holocentric, or their centromeres might be epigenetically defined (Khalaf et al., 2024; Mascarenhas dos Santos et al., 2023).

**Figure 4:**
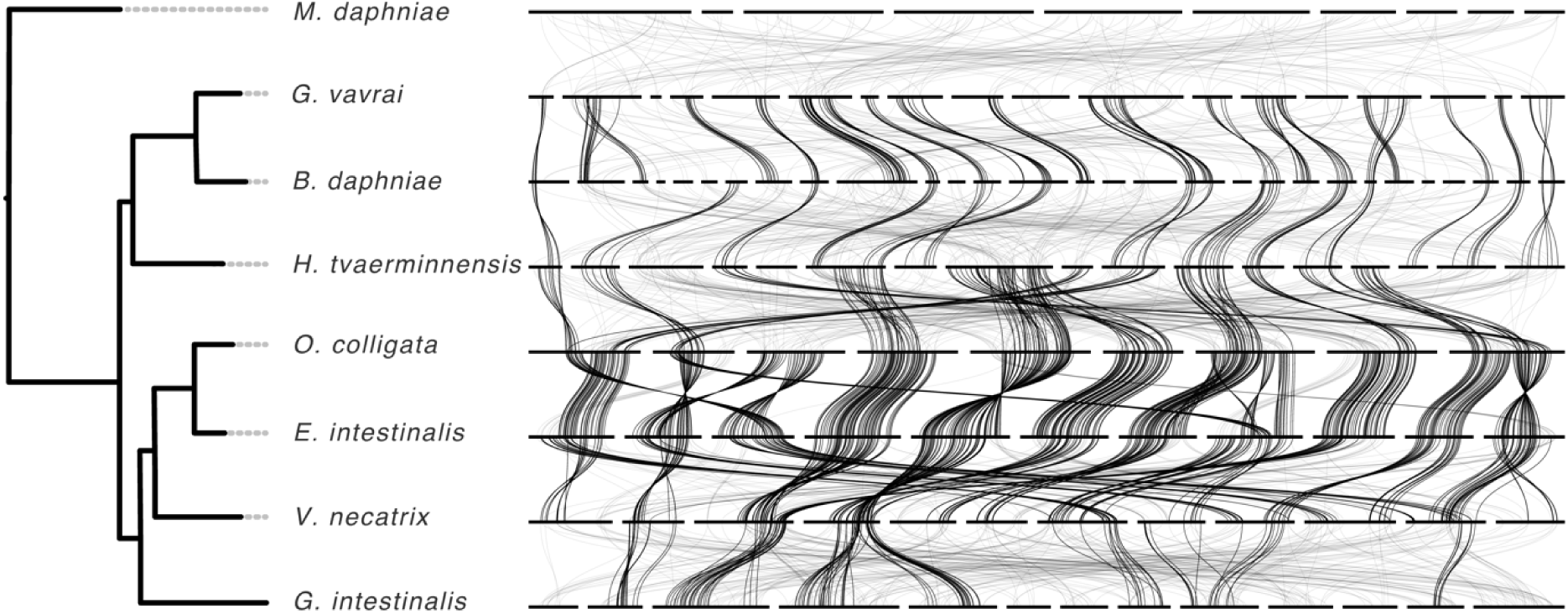
Ribbon diagram with maximum likelihood phylogeny showing synteny limited to closely related microsporidia. Contigs, some of which represent complete chromosomes, are represented by horizontal bars and connected by ribbons that connect the location of individual single-copy BUSCO genes across assemblies. Bold-type ribbons represent statistically conserved linkages. Because of its lower assembly contiguity, *Conglomerata obtusa* was omitted from the analysis.

### Repeat landscapes

*Encephalitozoon* spp. have few to no repeats except at chromosome ends, which are all highly similar (Mascarenhas dos Santos et al., 2023). However, as seen in Figure 1, the genomes analyzed here showed high variation in repeat content, suggesting that repeats may play a role in shaping the evolution of microsporidian genomes. Using the Kimura substitution level, which estimates divergence from a repeat consensus sequence, we assessed repeat content and its activity over evolutionary time. High substitution levels indicate high divergence related to activity deep in the evolutionary past, and low substitution levels indicate low divergence and therefore recent activity. The largest genomes showed high repeat content and constant repeat activities throughout their past (Figure 5). The three Amblyosporida species shared similar repeat activity, especially in their early past, indicative of the activity in a common ancestor. DNA and LTR transposons were highly active in all three species, but *G. vavrai*, and especially *B. daphniae*, showed higher activity in the recent past. The apparent lack of recent activity in *C. obtusa* might be an artefact of the low divergence repeats collapsed in this short-read assembly. Recent increases in repeats compared to earlier activity were more pronounced in species with smaller genomes, such as LINEs in *G. intestinalis.* These observations all indicate that different transposon families could shape the larger genomes. As species with shorter genomes had less activity in their more distant past, repeats may have only recently been expanded or removed in the shorter genomes.

**Figure 5:**
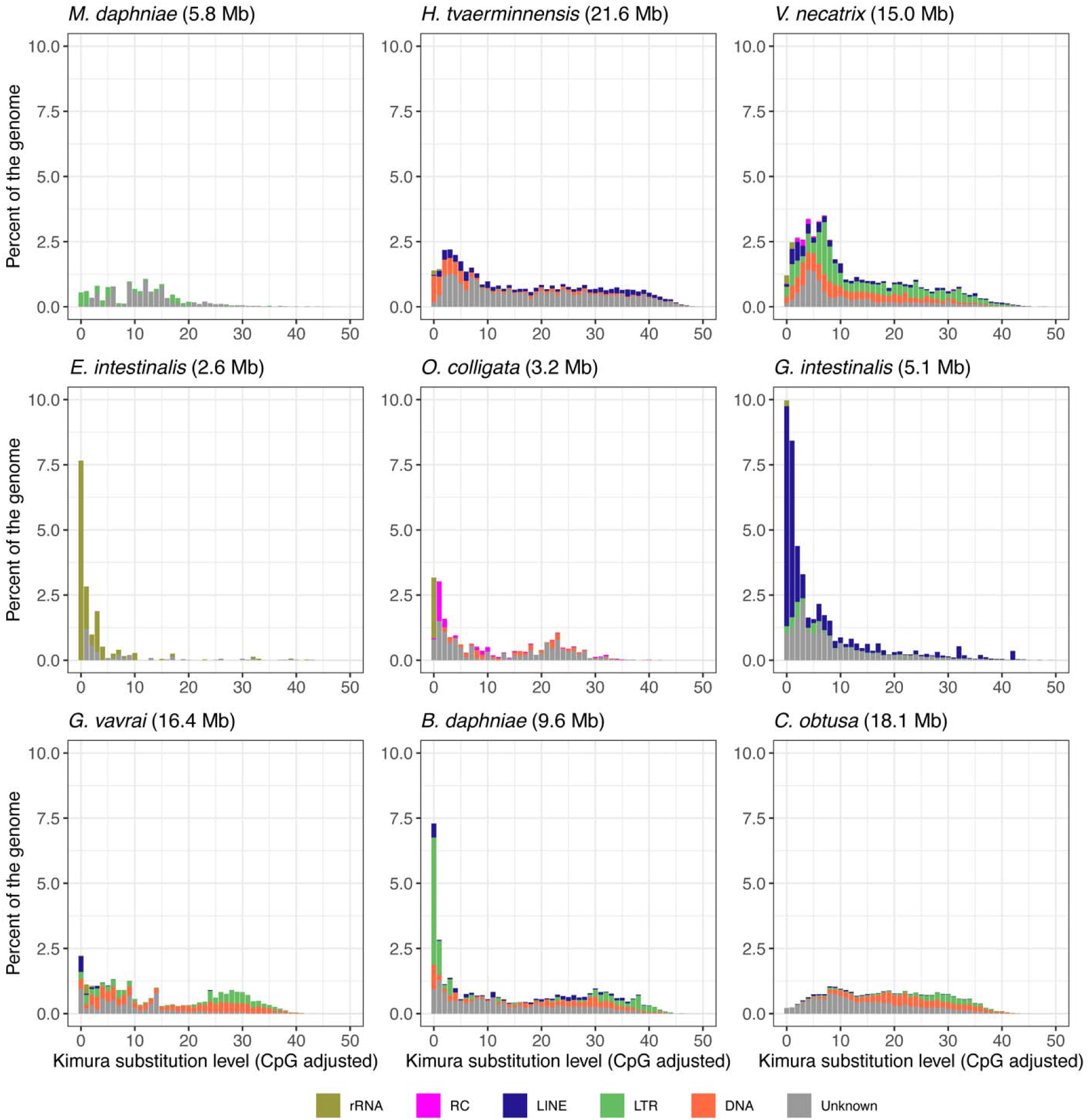
Repeat landscapes of microsporidia. Low Kimura substitution levels represent low divergence from the repeat consensus sequence, corresponding to recently expanded sequences, whereas high substitution levels indicate high divergence, indicating earlier activity. Genome assembly lengths are indicated in brackets.

### Methylation

Epigenetic modifications may act to regulate not only gene expression but also repeat element activity and the associated variation in genome architecture. *Encephalitozoon* has hypermethylated chromosome ends with rRNA genes and hypomethylated chromosome cores with mRNA genes (Mascarenhas dos Santos et al., 2023). Using a genome ontology approach based on hypergeometric tests, we compared these methylation patterns with the genomes for which we have epigenetic information. From our species with available ONT data for methylation analysis, we found that the closest *Encephalitozoon* relative, *G. intestinalis*, showed the most similar pattern: hypermethylated rRNA genes at the chromosome ends and hypomethylated mRNA genes at the chromosome cores (Tables 2 and S3). Unlike *Encephalitozoon*, however, *G. intestinalis* had LTRs and hypermethylated LINE repeats. In the other species with available ONT data, *H. tvaerminnensis*, *G. vavrai*, and *B. daphniae* also had hypermethylated rRNA genes and tended to have hypermethylated transposable repeat elements like DNA, LINE, and LTR, even though their genome architectures were very different than *Encephalitozoon* and *G. intestinalis*. Non-transposable repeats, like simple repeats and low-complexity regions, were hypomethylated in all species. Coding sequences were hypomethylated in the *H. tvaerminnensis* genome, like in *Encephalitozoon* and *G. intestinalis*, but were hypermethylated in *G. vavrai* and *B. daphniae*. Differential methylation of coding sequences and repeat elements could indicate gene expression and repeat activity regulation, respectively. The two repeat elements that expanded most recently (LINEs in *G. intestinalis* and LTRs in *B. daphniae*) were not consistently hyper- or hypomethylated.

**Table 2:** Methylation patterns of microsporidia. Overrepresented (= hypermethylated) genomic features have a negative log *P-*value, whereas underrepresented (= hypomethylated) genomic features have a positive log *P-*value. Asterisks denote levels of statistical significance obtained from genome ontology, while shading highlights directionality. Specifically, **\*** indicates p < 0.05, **\*\*** indicates p < 0.01, and **\*\*\*** indicates p < 0.001.

|  | *E. intestinalis* | *B. daphniae* | *G. vavrai* | *G. intestinalis* | *H. tvaerminnensis* |
| --- | --- | --- | --- | --- | --- |
| DNA repeat | — | 4.41 * | -21.86 *** | — | -202.4 *** |
| LINE repeat | — | 5.06 ** | 1.46 | -136.83 *** | -11.48 *** |
| Low complexity | 2.90 | 4.44 * | 21.08 *** | 1 | 13.75 *** |
| LTR | — | -1.91 | -91.96 *** | -1.58 | -7.53 *** |
| mRNA | 1,476.72 *** | -5.39 ** | -4.91 ** | 77.31 *** | 19.55 *** |
| RC repeat | — | — | — | — | -1.25 |
| rRNA | -2,936.43 *** | -72.86 *** | -35.97 *** | -107.06 *** | -1.05 |
| Satellite repeat | — | — | — | — | -2.73 |
| Simple repeat | 17.22 *** | 14.72 *** | 62.19 *** | 7.99 *** | 31.30 *** |
| Unknown repeat | -163.04 *** | -17.66 *** | -1.9 | -200.41 *** | -8 *** |

### Present and absent protein domains

Genome streamlining in microsporidia has resulted in the loss of protein domains that are considered essential in non-parasitic free-living fungi (Haag et al., 2014; Jespersen et al., 2022). Using comparative genomics, we identified present and absent metabolic and functional capacities across microsporidia, looking at which functions were retained in all or most species and which were reduced. We began by summarizing the presence and absence of protein domains (Pfam)–characteristic protein families for biological pathways (Table S4). Among the nine microsporidia genomes, the extended microsporidium *M. daphniae* had, as expected, the most exclusive protein domains (656), even more than the shared protein domains across all nine species (444). These included mitochondrial carrier proteins (PF00153) that are characteristic of the transfer of molecules across mitochondrial membranes (*M. daphniae* has a mitochondrion) but that were absent in all mitochondria-free canonical microsporidia (Figure 6). Asparyl protease domains, e.g., PF13650, were among the most abundant protein domains with presence/absence polymorphism among the nine microsporidia (Figure 6). These endopeptidases are known virulence factors of pathogenic fungi (Mandujano-González et al., 2016). Looking at all species used in our phylogeny (Figure 2), we found that the functional glycolysis characteristic Fructose-bisphosphate aldolase class-I family (PF00274) was absent in *Enterocytozoon bieneusi* (Table S4), consistent with previous findings that glycolysis is reduced in this species that obtains its energy directly from the host (Wiredu Boakye et al., 2017). Protein domains like ABC transporter (PF00005), Hsp70 (PF00012), and actin (PF00022) were present in all species (Table S4) and thus appear to be essential for microsporidia.

**Figure 6:**
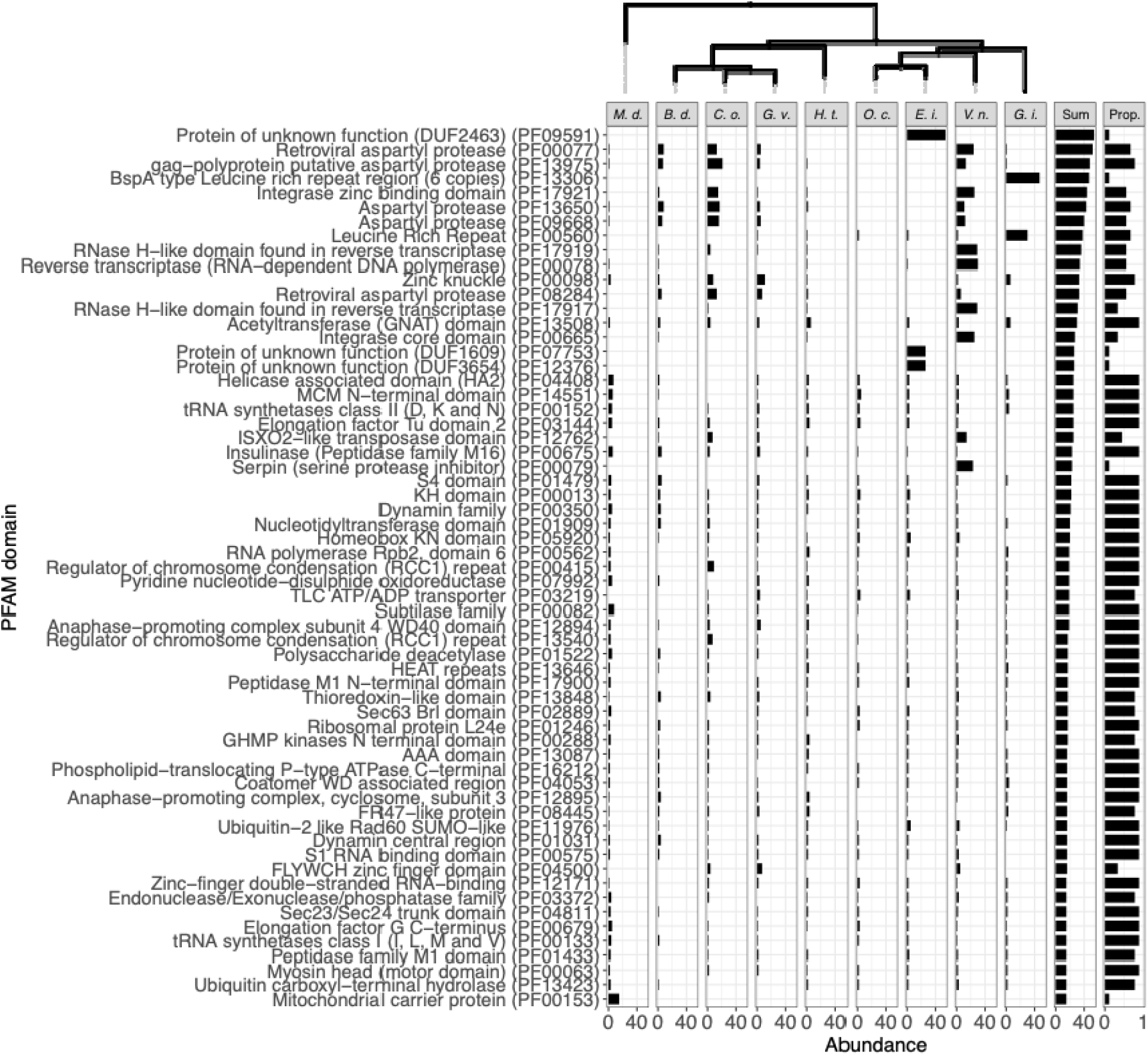
Variably present protein domains (Pfam) ordered from most to least abundant. Rows list individual protein domains with their Pfam domain description and ID (in parentheses), while columns with a maximum likelihood phylogeny at the top list the nine species. The penultimate column tallies the number of protein domains present, while the last column provides the proportion of species in which they occur. Full data is presented in Table S4.

## Discussion

Microsporidia are intracellular parasites that are enigmatic due to their high degree of specialization. Our goal was to better understand the evolution of their genome by producing several new high-quality genome assemblies that reveal the (epi)genetic architecture across this taxon. We focused mainly on defining which features are general for all microsporidia and which are derived in subclades. GC-content, genome length, chromosomal sequence arrangement, and methylation patterns vary widely among microsporidian clades; however, species that share a more recent common ancestor tend to exhibit less variation in these features, similar to other systems with high-quality genomes available for study. The architecture of microsporidia genomes studied here were only similar to the previously described *Encephalitozoon* in one close relative and in a species of the next closest major microsporidian clade, proving that *Encephalitozoon*’s chromosomal sequence arrangement is not general to all microsporidia, as has been speculated (Brugère et al., 2000). Our other studied species showed very diverse chromosomal arrangements and repeat contents, although the same genetic features tend to be hyper- or hypomethylated in all species. For example, rRNA gene positions are different among all microsporidian clades, but always hypermethylated in all species. Because the new and improved microsporidia genome assemblies presented here are all from species that infect the aquatic crustacean *Daphnia,* we can see that the variation in genome architecture is not explained by the host: indeed, the host’s role in parasite genome architecture appears to be minor. As models, thus, *Daphnia* and its microsporidian parasites from across the phylum allow us to extend microsporidia research and improve our understanding of genome evolution in these extreme parasites beyond the mammal-parasite model.

### Nucleotide composition linked to genome size

Nucleotide composition is fundamental to genome architecture, and although GC-content varies widely across all lifeforms, it tends to evolve to lower values in parasites such as the microsporidia. Reduced recombination rates, diminished DNA repair mechanisms, relaxed purifying selection, and increased selection for AT mutations have all been proposed as reasons for reduced GC in parasite genomes (Videvall, 2018). These mechanisms may play a role in the negative correlation observed here between GC-content and repeat content, which is also seen in fungi and other related phyla (Elliott & Gregory, 2015). First, it might be easier for repeat elements to proliferate in species with relaxed purifying selection due to the small effective population size relative to the mode of transmission (Haag et al., 2020). We expect species with a small effective population size to have more and more recent expansions of repeat elements due to vertical transmission compared to species with a higher effective population size and horizontal-only transmission (de Albuquerque et al., 2020). With our sample, it is hard to draw conclusions on demographic effects; however, long genome Amblyosporida and Nosema/Vairimorpha clade species are known for complex life cycles with mixed-mode transmission (Vávra et al., 2018; Xiong et al., 2023), and mixed-mode transmission in the large genome microsporidium *H. tvaerminnensis* has been associated with small effective population sizes (Haag et al., 2020). Contrarily, species with small genomes are only transmitted horizontally. Second, AT mutations might accumulate in proliferating repeat elements, directly and indirectly accelerating the decrease in GC-content. This occurs indirectly if 5mC methylation is used to slow the proliferation of repeat elements because fewer positions are available for this epigenetic modification. In contrast, GC-richer repeats might be more of a methylation target, possibly constraining their proliferation. Although repeat elements were hypermethylated across species, repeat activity was more evenly distributed across time in lower GC species than in higher GC species, where activity occurred mainly in the recent past. This could be because higher GC species have more stringent selection against repeat proliferation, more efficient mechanisms to remove repeats, higher recombination rates, or lower tendency for AT mutation, such that our methodology estimates higher recent repeat activity because of the lower divergence among them.

### Distribution of rRNA genes

The rRNA genes are of special interest in microsporidia genome architecture, mainly because of their unusual location in *Encephalitozoon* close to each chromosome end, at the subtelomeres (Mascarenhas dos Santos et al., 2023). Contrary to previous findings from a short-read-based draft genome (Mascarenhas dos Santos et al., 2023; Pombert et al., 2015), we show that *O. colligata* shares this structure with its close *Encephalitozoon* relatives. This highlights the use of high-quality resources to study genome architecture. rRNA genes are also located at some, but not all, of the subtelomeres in a species of the next closely related major clade, *G. intestinalis,* which also shows small-scale rRNA gene rearrangements (Refardt & Mouton, 2007). Given that transposable elements (TEs) show high and recent activity in *G. intestinalis*, that rRNA genes and TEs are co-located, and that TEs interact with and benefit from each other (Garcia et al., 2024), we suggest that rRNA genes and TEs may coevolve together in this microsporidium or even across microsporidia, a hypothesis supported also by our finding that rRNA genes and TEs are co-located in *H. tvaerminnensis* and Amblyosporida microsporidia. Moreover, rRNA genes in these species have yet another outstanding arrangement: rRNA operons occur in tandem with tail-to-tail orientation and few LINE or unknown repeats between them. These rRNA operon pairs are not located close to a chromosome end, however, suggesting that the proximity of rRNA genes to chromosome ends is a derived characteristic found in some species of the Nosematida and Enterozytozoonida clades.

### Co-occurrence and epigenetic modification of repeat elements

The significance of methylation is unclear in microsporidia. Mascarenhas dos Santos et al. (2023) have speculated that methylation at rRNA genes facilitates heterochromatin formation, helping to silence the ribosomal machinery during phases of the life cycle that have low or no access to energy. Indeed, we found rRNA genes to be hypermethylated in all the species studied, regardless of their location. The hypothesis proposed by Mascarenhas dos Santos et al. (2023) could also apply to other genetic loci, such as TEs, which because their activity may be unfavorable, are silenced by epigenetic modifications that facilitate heterochromatin formation. Whereas small microsporidia genomes have low to no repeats, including TEs, and hypomethylated cores, larger genomes might show methylation at repeat loci across the genome to reduce TE activity through heterochromatin formation. Given that rRNA genes and transposable elements co-occur in the studied microsporidia, hypermethylation could help to simultaneously control their activity and spread. At the same time, their co-occurrence might be beneficial to the TEs, which might be more likely to proliferate or less likely to be removed if they occur close to the rRNA genes, which are highly expressed. The microsporidia *V. necatrix* and *Nosema muscidifuracis* from the same clade as *Encephalitozoon* and *Ordospora* have more randomly distributed rRNA operons (and *N. muscidifuracis* has many more), while both have much lower whole-genome GC-content than their relatives (Xiong et al., 2023). A telomere-to-telomere assembly of the *N. muscidifuracis* genome would help pin down rRNA gene location and methylation data for the two species to clarify the role of epigenetic modification in the evolution of microsporidia repeat families. Neither epigenetic, nucleotide, nor sequence composition has indicated the location of centromeres in the species studied here. Hi-C sequencing could help identify centromeres (Khalaf et al., 2024).

## Conclusion

Microsporidia have attracted much attention because of their parasitic life history and because they include some of the smallest known eukaryote genomes. However, their genomes also exhibit drastic variation in size and other features. Here we present high-quality genome assemblies from across the major microsporidia clades, showing covariance in GC-content, chromosomal sequence arrangement, and epigenetic modification. Species with high GC-content have a sequence arrangement similar to *Encephailtozoon* spp., with hypermethylated subtelomeres and hypomethylated cores. In contrast, low-GC species do not share this pattern, having instead a higher activity of repeat elements, which have lower GC-content than coding sequences, likely because the latter are under purifying selection (Videvall, 2018). Low-GC genomes might arise due to a lowered selection regime that allow transposable repeats to proliferate more easily (de Albuquerque et al., 2020; Videvall, 2018). The evolutionary dynamics leading to the discrepancy between relatively strong selection on coding sequences and weak selection outside of them remain unknown. Still, the activity of repeat elements clearly shapes the evolution of genome architecture, and with the recent advent of high-throughput sequencing technologies should thus be included in comparative studies of GC-content where it was previously not.

Although the streamlining of genomes has led to a significantly fewer coding sequences across all core microsporidia (Žárský et al., 2023), certain species show extraordinary genome reductions, like the partial absence of glycolysis genes in *Enterocytozoon* spp. (Wiredu Boakye et al., 2017). Here, we provide an overview of the presence and absence of protein domains across microsporidia as a resource for studies focused on the presence/absence polymorphism and functional adaptations to parasitism across the microsporidia clades. Knowing the phylogenetic distribution of protein domains and other genomic features in the microsporidia allows us to dive deeper into the evolution of this extraordinary group of parasites. Features that are conserved by all microsporidia may have contributed to the radiation of this group, with the loss of typical mitochondria being the hallmark example. In contrast, features present only in individual microsporidian clades or species may have been important in evolving adaptations to specific hosts or life histories. Furthermore, correlations between the presence of certain genomic features and life history traits can shed light on the evolutionary drivers of these traits. Thus, the presence or absence of protein domains and other genomic features can be the starting point for in-depth comparative study of the microsporidia and their genomic architecture with these high-quality genomes. As a final step for obtaining telomere-to-telomere assemblies, especially the harder-to-assemble longer genomes, we suggest applying Hi-C sequencing to microsporidia.

## Methods

### Samples

Using long wide-mouth pipettes, we collected *Daphnia pulex* infected with *C. obtusa* from the Tvaerminne archipelago, Finland (59°49’55.4", N 23°15’17.6" E), placed them in RNAlater (Ambion, Glasgow, United Kingdom), and kept them refrigerated until DNA isolation. *Daphnia pulex* infected with *G. vavrai* were collected from Aegelsee lake near Frauenfeld, Switzerland (47°33′28.0″ N, 8°51′46.0″ E) and were transported to the laboratory for DNA isolation. *Daphnia magna* infected with *B. daphniae* (ID: BE-OM-3), *G. intestinalis* (KZ-23-1), and *O. colligata* (GB-LK1-1) were previously collected and kept in the lab as iso-female lines (Angst et al., in prep.; Pombert et al., 2015; Refardt et al., 2008) until DNA isolation. We obtained high-molecular-weight DNA from *Daphnia* with parasites using the GenePure DNA Isolation Kit (QIAGEN, Hilden, Germany) as described in Angst et al. (in review) (https://dx.doi.org/10.17504/protocols.io.5jyl82n96l2w/v1; Angst & Fields (2024)). Illumina paired-end sequencing and either PacBio HiFi (Pacific Biosciences high fidelity) or ONT (Oxford Nanopore Technologies) sequencing was used to obtain the highest quality nucleotide or modification calls, respectively. Illumina libraries were prepared using Kapa PCR-free kits and sequenced by the Quantitative Genomics Facility service platform at the Department of Biosystem Science and Engineering (D-BSSE, ETH) Basel, Switzerland, on an Illumina HiSeq 6000 sequencer. PacBio SMRTbell libraries were prepared and sequenced on a PacBio Revio at the Lausanne Genomics Technologies Facility (GTF, UNIL), Switzerland. We prepared ONT libraries with the SQK-LSK110 ligation kit and sequenced them using a MinION device with a Spot-ON Flow Cell (R9.4.1). Additionally, we downloaded genomic data of *E. intestinalis* (NCBI database; Assembly name: ASM2439929v1; GenBank assembly accession: GCA_024399295.1; SRA accession: SRR17865590; Bioproject accession: PRJNA594722; Mascarenhas dos Santos et al. 2023) *H. tvaerminnensis* (NCBI database; Assembly name: FIOER33 v3; GenBank assembly accession: GCA_022605425.2; SRA accession: SRR24575619, Bioproject accession: PRJNA778105; Angst et al. 2023), *M. daphniae* (NCBI database; Bioproject accession: PRJNA1199805; Angst et al. (in prep.)), and *V. necatrix* (NCBI database; Assembly name: ASM3663032v1; GenBank assembly accession: GCF_036630325.1; Bioproject accession: PRJNA909071; Svedberg et al. 2024) for analysis.

## Assembly

### Glugoides intestinalis *and* O. colligata

*Glugoides intestinalis* (total/average read length: 79.38 Gb/10.75 Kb) and *O. colligata* (20.22 Gb/8.84 Kb) were sequenced using PacBio HiFi sequencing to an average sequencing coverage of 55× and 45×, respectively. We used hifiasm v.0.19.8-r603 (Cheng et al., 2021) to assemble the obtained sequencing reads with default parameters for *G. intestinalis* and with --hom-cov 45 -l1 --n-hap 1 parameters for *O. colligata*. Additionally, we generated a second hifiasm assembly for both species after excluding host-specific PacBio HiFi reads using SAMtools v.1.7 (Danecek et al., 2021) that were identified by mapping them to the *D. magna* genome (NCBI database; Assembly name: ASM4014379v1; GenBank assembly accession: GCA_040143795.1; BioProject ID: PRJNA624896; Cornetti et al. 2024) using minimap2 v.2.20-r1061 (H. Li, 2018). For each species, we aligned the two assemblies using minimap2 and merged them manually. We assessed the quality and completeness of the assemblies using QUAST v.5.0.2 (Gurevich et al., 2013) and BUSCO v.5.5.0 (Manni et al., 2021) with the microsporidia_odb10 database (Creation date: 2020-08-05). All software was used with default parameters unless stated otherwise.

### Binucleata daphniae *and* G. vavrai

We sequenced *B. daphniae* (total/N50: 0.72 Gb/7.21 Kb) and *G. vavrai* (3.96 Gb/3.53 Kb) using ONT sequencing in adaptive mode, depleting for sequences homologous to the *D. magna* and the *D. pulex* genome (NCBI database; Assembly name: PA42 4.2; GenBank assembly accession: GCA_911175335.1; BioProject ID: PRJEB46221; Ye et al. 2021), respectively. We used bonito v.0.5.3 (https://github.com/nanoporetech/bonito) to base-call ONT sequencing reads with the dna_r9.4.1_e8.1_sup@v3.3 model. For the *B. daphniae* genome assembly, we used MaSuRCA v.4.0.5 (Zimin et al., 2017) with ONT (53×) and Illumina (639×) sequencing reads, excluding host-specific, adaptor, and low-quality sequences using minimap2, bwa-mem2 mem v.2.2.1 (Vasimuddin et al., 2019), SAMtools, and fastp v.0.23.2 (Chen, 2023) with -q 30 and -l 100 parameters. Haplotigs were purged twice from the assembly using the Purge Haplotigs pipeline v.1.1.2 (Roach et al., 2018) with -l 125 -m 220 -h 400 parameters and BEDTools v.2.30.0 (Quinlan, 2014) for generating the read-depth distribution. For the *G. vavrai* genome assembly, we used nextDenovo v.2.5.0 (Hu et al., 2024) with ONT sequencing reads (98×) and polished the resulting assembly with Illumina sequencing reads (458×) using nextPolish v.1.4.1 (Hu et al., 2020), Medaka v.1.7.2 (https://github.com/nanoporetech/medaka) and Pilon v.1.24 (Walker et al., 2014). The Purge Haplotigs pipeline was applied with -l 88 -m 176 -h 300 parameters.

### Conglomerata obtusa

Because we were unable to generate sufficient quality genomic material to conduct long-read sequencing for *C. obtusa*, we used Illumina sequencing (153×) for this case only. We excluded sequencing reads mapping to the *D. pulex* host genome and assembled the remaining reads using megahit v.1.2.9 (D. Li et al., 2016). We excluded contigs with less than a third of the average coverage, shorter than 500 basepairs, with GC-content above 33, or with coverage over 1× for a *Daphnia* sample from a parasite-free pond of the same island (NCBI database; SRA accession: SRR26394761, Bioproject ID: PRJNA862292; Angst et al. 2024) using seqtk v.1.3 (https://github.com/lh3/seqtk) and BBMap v.38.96 (Bushnell, 2014).

### Genome annotation and phylogenetic inference

We annotated *de novo* genome assemblies as described in Angst et al. (in prep.) using funannotate v.1.8.14 (Palmer & Stajich, 2022) and associated software for gene prediction and functional annotation (Almagro Armenteros et al., 2019; Brůna et al., 2020; Cantalapiedra et al., 2021; Frith, 2011; Haas et al., 2008; Huerta-Cepas et al., 2019; Jones et al., 2014; Käll et al., 2004; Korf, 2004; Majoros et al., 2004; Seppey et al., 2019; Stanke et al., 2006, 2008). For phylogenetic inference, we downloaded genomes from a broad range of microsporidia (Aurrecoechea et al., 2017) including the extended microsporidium *P. saccamoebae* (Quandt et al., 2017) along with *Rozella allomycis* (James et al., 2013) as an outgroup. We identified single-copy orthologs among the species using proteinortho v.6.3.0 (Klemm et al., 2023), concatenated them, aligned the individual genes as well as the concatenated genes using MAFFT v.7.508 (Katoh et al., 2002; Katoh & Standley, 2013), and trimmed the alignments using trimAl v.1.4.rev15 (Capella-Gutiérrez et al., 2009) with the -automated1 flag. We inferred a maximum likelihood species tree from the concatenated sequences using IQ-TREE 2 v.2.2.0 (Minh et al., 2020) and a weighted phylogeny from the individual maximum likelihood gene trees using Weighted ASTRAL v.1.19.3.7 (Zhang & Mirarab, 2022) with - r 16 -s 16 -x 100 -n 0 parameters separately for all species and the nine main study species.

### Comparative genomics and genome ontology

We looked for homology and synteny across genomes using odp v.0.3.3 (Schultz et al., 2023). Specifically, we used the coordinates of the identified BUSCO genes to infer ancestral BUSCO gene linkage groups, test for conserved linkages among genomes, and create the ribbon diagram. To obtain modified base calls from ONT sequencing reads, we used megalodon v.2.5.0 (https://nanoporetech.github.io/megalodon) focusing on 5mC modifications for downstream analyses. We tested whether genomic annotations, including mRNA genes, rRNA genes, and different repeat types, were enriched with methylation sites by using the genome ontology peak annotation enrichment in homer v.5.1 (Heinz et al., 2010). This analysis considered sites to be methylated if read depth was above 3× and percent modified reads over 50 %. The inclusion of multiple cell stages in our DNA isolation protocol explains, at least in part, variation in epigenetic modification between sequencing reads. For example, spores might have a different methylation pattern than sporonts, meronts, or other life stages (Mascarenhas dos Santos et al., 2023). For *G. intestinalis*, we re-used ONT reads from Angst et al. (in prep.) (NCBI database; SRA accession: SRR31799445, Bioproject accession: PRJNA1199805). We identified repeats using RepeatModeler v.2.0.2, including the LTR pipeline (Flynn et al., 2020), and ReapeatMasker v.4.1.2 (Smit et al., 2013), and separately identified rRNA genes using cmscan of the Infernal pipeline v.1.1.2 (Nawrocki & Eddy, 2013). From the ReapeatMasker alignments, we created repeat landscapes using ReapeatMasker utils calcDivergenceFromAlign.pl and createRepeatLandscape.pl. Finally, we summarized PFAM domains to identify the presence and absence of protein domains across all species using the module compare of funannotate, and the tidyverse 2.0.0 (Wickham et al., 2019) R v.4.4.1 (R Core Team, 2024) package for representation and statistics.

## Data availability

Raw data are deposited in the NCBI SRA database, and the assembled genome plus the predicted protein sequences are available in the NCBI GenBank (BioProject ID: PRJNA1199946).

## Acknowledgments

We thank Ju rgen Hottinger, Michelle Krebs, and Urs Stiefel for their help in the laboratory and all members of the Ebert group for providing feedback on the study and the manuscript. Alix Thivolle extracted DNA from *D. pulex* infected with *C. obtusa*. Christina Tadiri collected *D. pulex* infected with *G. vavrai*. Calculations were performed at sciCORE (http://scicore.unibas.ch/) scientific computing center at the University of Basel. Suzanne Zweizig improved the language of the text.

