## Supplementary material for "Variation in genome architecture and epigenetic modification across the microsporidia phylogeny": Figures S1 and S2 and Tables S1 to S3

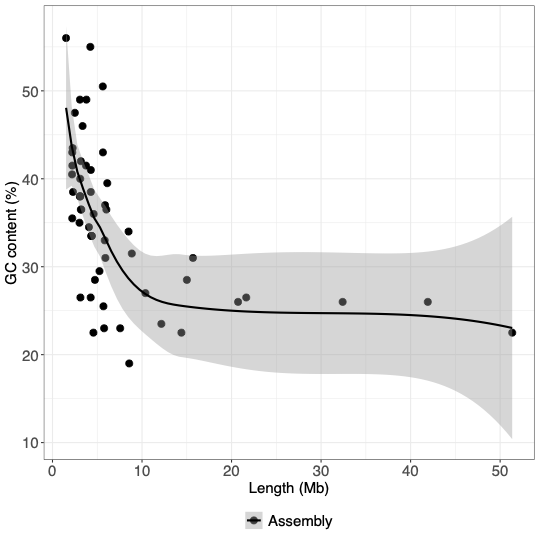

**Figure S1: Relationship between genome assembly length and GC-content for all available NCBI microsporidia reference genomes.** The line represents local regressions with LOESS; the 95% confidence interval is indicated in gray.

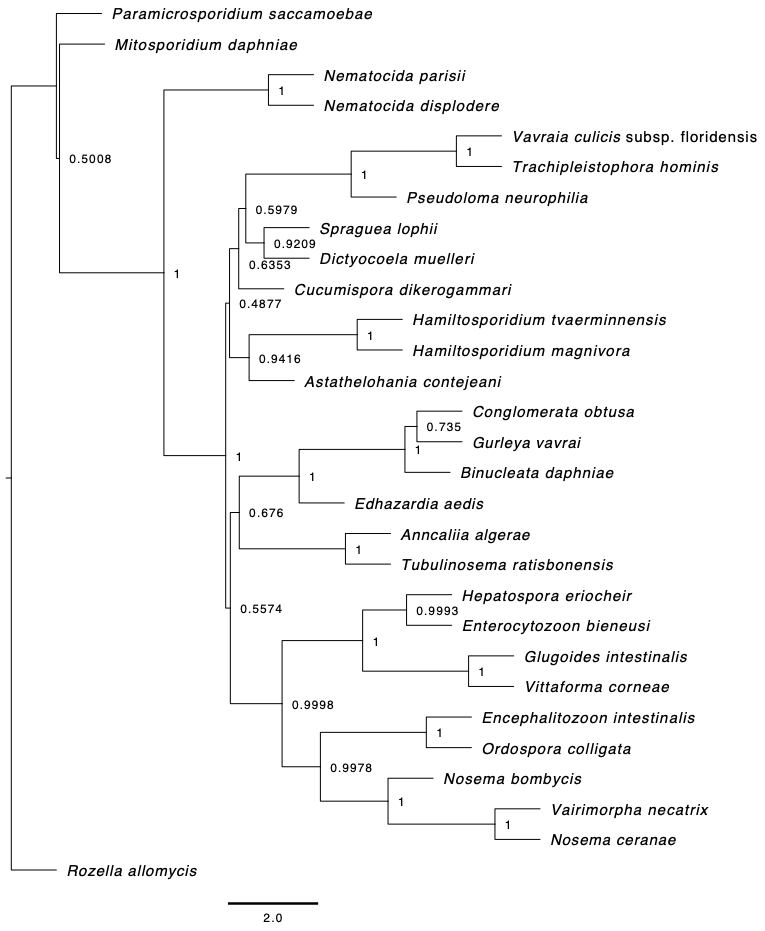

**Figure S2: Weighted phylogeny from individual maximum likelihood gene trees using Weighted ASTRAL.** The species tree obtained by maximum likelihood estimation from concatenated gene sequences (Figure 2) is more similar, topologically, to the weighted phylogeny among closely related species than across the major clades. Note that terminal branch lengths are not calculated by ASTRAL. Unlike previous rRNA gene phylogenies, where *C. obtusa* has been the outgroup to *B. daphniae* and *G. vavrai*, our phylogenetic species tree shows *B. daphniae* as the outgroup within the Amblyosporida*.*

**Table S1: Microsporidia reference genome assemblies in NCBI GenBank with length and GC-content.**

| **GanBank assembly accession** | **Scientific name** | **Assembly length (Mb)** | **GC-content** |
| --- | --- | --- | --- |
| GCA_036630325.1 | *Vairimorpha necatrix* | 15.01 | 28.5 |
| GCA_000091225.2 | *Encephalitozoon cuniculi* GB-M1 | 2.498 | 47.5 |
| GCA_000146465.1 | *Encephalitozoon intestinalis* ATCC 50506 | 2.217 | 41.5 |
| GCA_022605425.2 | *Hamiltosporidium tvaerminnensis* | 21.64 | 26.5 |
| GCA_000277815.3 | *Encephalitozoon hellem* ATCC 50504 | 2.252 | 43.5 |
| GCA_000280035.2 | *Encephalitozoon romaleae* SJ-2008 | 2.188 | 40.5 |
| GCA_007674295.1 | *Antonospora locustae* | 3.171 | 42 |
| GCA_021821965.1 | *Ordospora pajunii* | 2.206 | 43 |
| GCA_000760515.2 | *Mitosporidium daphniae* | 5.635 | 43 |
| GCA_021653875.1 | *Nematocida major* | 3.787 | 49 |
| GCA_024244105.1 | *Nematocida minor* | 3.742 | 41.5 |
| GCA_024243835.1 | *Enteropsectra breve* | 6.016 | 36.5 |
| GCA_026692725.1 | *Enterocytozoon hepatopenaei* | 3.119 | 26.5 |
| GCA_000988165.1 | *Vairimorpha ceranae* | 5.691 | 25.5 |
| GCA_000383075.1 | *Nosema bombycis* CQ1 | 15.69 | 31 |
| GCA_036780745.1 | *Vairimorpha bombi* | 4.739 | 28.5 |
| GCA_001887945.1 | *Spraguea lophii* | 5.765 | 23 |
| GCA_001642395.1 | *Nematocida displodere* | 3.066 | 49 |
| GCA_000803265.1 | *Ordospora colligata* OC4 | 2.291 | 38.5 |
| GCA_000738915.1 | *Nematocida ausubeli* | 4.276 | 38.5 |
| GCA_000250985.1 | *Nematocida parisii* ERTm1 | 4.071 | 34.5 |
| GCA_000192795.1 | *Vavraia culicis* subsp. floridensis | 6.119 | 39.5 |
| GCA_000231115.1 | *Vittaforma corneae* ATCC 50505 | 3.214 | 36.5 |
| GCA_000447185.1 | *Vairimorpha apis* BRL 01 | 8.57 | 19 |
| GCA_000316135.1 | *Trachipleistophora hominis* | 8.498 | 34 |
| GCA_001432165.1 | *Pseudoloma neurophilia* | 5.248 | 29.5 |
| GCA_002087885.1 | *Hepatospora eriocheir* | 4.57 | 22.5 |
| GCA_000230595.3 | *Edhazardia aedis* USNM 41457 | 51.35 | 22.5 |
| GCA_000385875.2 | *Anncaliia algerae* PRA339 | 12.16 | 23.5 |
| GCA_038659295.1 | *Enterocytospora artemiae* | 3.015 | 35 |
| GCA_028335825.1 | *Nosema muscidifuracis* | 14.4 | 22.5 |
| GCA_024244045.1 | *Nematocida* sp. AWRm80 | 3.172 | 36.5 |
| GCA_024243985.1 | *Nematocida* sp. LUAm3 | 3.092 | 38 |
| GCA_024244025.1 | *Nematocida* sp. LUAm2 | 3.063 | 38 |
| GCA_024244035.1 | *Nematocida* sp. LUAm1 | 3.081 | 38 |
| GCA_043388365.1 | *Albopleistophora grylli* | 5.874 | 37 |
| GCA_024244115.1 | *Nematocida* sp. AWRm77 | 3.362 | 46 |
| GCA_001642415.1 | *Nematocida* sp. ERTm5 | 4.398 | 33.5 |
| GCA_029232285.1 | *Agmasoma* sp. JBojko-2022a | 4.265 | 26.5 |
| GCA_024244095.1 | *Nematocida homosporus* | 4.282 | 41 |
| GCA_024243935.1 | *Nematocida* sp. AWRm79 | 4.349 | 33.5 |
| GCA_024244055.1 | *Nematocida* sp. AWRm78 | 4.331 | 33.5 |
| GCA_023647555.1 | *Pancytospora philotis* | 4.227 | 55 |
| GCA_014805555.1 | *Astathelohania contejeani* | 10.38 | 27 |
| GCA_002087915.1 | *Enterospora canceri* | 3.095 | 40 |
| GCA_003600395.1 | *Metchnikovella incurvata* | 5.835 | 33 |
| GCA_004000155.1 | *Tubulinosema ratisbonensis* | 7.56 | 23 |
| GCA_001875675.1 | *Amphiamblys* sp. WSBS2006 | 5.62 | 50.5 |
| GCA_004325065.1 | *Hamiltosporidium magnivora* | 20.73 | 26 |
| GCA_015832245.1 | *Nosema granulosis* | 8.86 | 31.5 |
| GCA_014805705.1 | *Cucumispora dikerogammari* | 32.41 | 26 |
| GCA_024243955.1 | *Pancytospora epiphaga* | 4.595 | 36 |
| GCA_016256075.1 | *Dictyocoela muelleri* | 41.92 | 26 |
| GCA_032191455.1 | *Microsporidia* sp. MB | 5.909 | 31 |
| GCA_030386735.1 | *Alternosema astaquatica* | 1.517 | 56 |
| GCA_016255985.1 | *Dictyocoela roeselum* | 2.206 | 35.5 |

**Table S2: rRNA gene and telomere locations in high-quality microsporidia genomes.** SSU = Small subunit; LSU = Large subunit.

| **Genomic feature per species** | **Contig_ID** | **Contig length (bp)** | **Feature start (bp)** | **Feature end (bp)** | **Stranded-ness** |
| --- | --- | --- | --- | --- | --- |
| ***B. daphniae*** | |  |  |  |  |
| SSU | scaffold_1 | 989,240 | 651,647 | 652,968 | + |
| LSU | scaffold_1 | 989,240 | 653,100 | 655,275 | + |
| LSU | scaffold_3 | 562,890 | 1,555 | 1 | - |
| SSU | scaffold_3 | 562,890 | 3,008 | 1,687 | - |
| SSU | scaffold_9 | 332,081 | 681 | 1 | - |
| SSU | scaffold_10 | 325,139 | 38,435 | 39,756 | + |
| LSU | scaffold_10 | 325,139 | 39,888 | 42,398 | + |
| telomere | scaffold_28 | 146,058 | at start |  | - |
| LSU | scaffold_29 | 125,248 | 125,248 | 124,516 | - |
| ***E. intestinalis*** |  |  |  |  |  |
| telomere | CP075158.1 | 189,841 | at start |  | - |
| LSU | CP075158.1 | 189,841 | 6,593 | 4,214 | - |
| SSU | CP075158.1 | 189,841 | 7,997 | 6,703 | - |
| SSU | CP075158.1 | 189,841 | 181,638 | 182,932 | + |
| LSU | CP075158.1 | 189,841 | 183,042 | 185,423 | + |
| telomere | CP075158.1 | 189,841 |  | at end | + |
| telomere | CP075159.1 | 213,886 | at start |  | - |
| LSU | CP075159.1 | 213,886 | 6,842 | 4,468 | - |
| SSU | CP075159.1 | 213,886 | 8,245 | 6,952 | - |
| SSU | CP075159.1 | 213,886 | 205,459 | 206,752 | + |
| LSU | CP075159.1 | 213,886 | 206,862 | 209,237 | + |
| telomere | CP075159.1 | 213,886 |  | at end | + |
| telomere | CP075160.1 | 217,008 | at start |  | - |
| LSU | CP075160.1 | 217,008 | 7,338 | 4,960 | - |
| SSU | CP075160.1 | 217,008 | 8,740 | 7,447 | - |
| SSU | CP075160.1 | 217,008 | 208,951 | 210,245 | + |
| LSU | CP075160.1 | 217,008 | 210,355 | 212,733 | + |
| telomere | CP075160.1 | 217,008 |  | at end | + |
| telomere | CP075161.1 | 227,972 | at start |  | - |
| LSU | CP075161.1 | 227,972 | 7,116 | 4,741 | - |
| SSU | CP075161.1 | 227,972 | 8,519 | 7,226 | - |
| SSU | CP075161.1 | 227,972 | 219,541 | 220,834 | + |
| LSU | CP075161.1 | 227,972 | 220,946 | 223,318 | + |
| telomere | CP075161.1 | 227,972 |  | at end | + |
| telomere | CP075162.1 | 226,151 | at start |  | - |
| LSU | CP075162.1 | 226,151 | 6,916 | 4,541 | - |
| SSU | CP075162.1 | 226,151 | 8,315 | 7,024 | - |
| SSU | CP075162.1 | 226,151 | 217,951 | 219,244 | + |
| LSU | CP075162.1 | 226,151 | 219,354 | 221,734 | + |
| telomere | CP075162.1 | 226,151 |  | at end | + |
| telomere | CP075163.1 | 230,490 | at start |  | - |
| LSU | CP075163.1 | 230,490 | 6,677 | 4,300 | - |
| SSU | CP075163.1 | 230,490 | 8,079 | 6,786 | - |
| SSU | CP075163.1 | 230,490 | 222,375 | 223,668 | + |
| LSU | CP075163.1 | 230,490 | 223,777 | 226,154 | + |
| telomere | CP075163.1 | 230,490 |  | at end | + |
| telomere | CP075164.1 | 239,752 | at start |  | - |
| LSU | CP075164.1 | 239,752 | 6,900 | 4,523 | - |
| SSU | CP075164.1 | 239,752 | 8,304 | 7,010 | - |
| SSU | CP075164.1 | 239,752 | 231,389 | 232,680 | + |
| LSU | CP075164.1 | 239,752 | 232,789 | 235,168 | + |
| telomere | CP075164.1 | 239,752 |  | at end | + |
| telomere | CP075165.1 | 247,125 | at start |  | - |
| LSU | CP075165.1 | 247,125 | 6,937 | 4,558 | - |
| SSU | CP075165.1 | 247,125 | 8,341 | 7,047 | - |
| SSU | CP075165.1 | 247,125 | 238,768 | 240,053 | + |
| LSU | CP075165.1 | 247,125 | 240,163 | 242,522 | + |
| telomere | CP075165.1 | 247,125 |  | at end | + |
| telomere | CP075166.1 | 268,753 | at start |  | - |
| LSU | CP075166.1 | 268,753 | 6,894 | 4,514 | - |
| SSU | CP075166.1 | 268,753 | 8,298 | 7,004 | - |
| SSU | CP075166.1 | 268,753 | 260,919 | 262,213 | + |
| LSU | CP075166.1 | 268,753 | 262,321 | 264,698 | + |
| telomere | CP075166.1 | 268,753 |  | at end | + |
| telomere | CP075167.1 | 277,983 | at start |  | - |
| LSU | CP075167.1 | 277,983 | 6,768 | 4,389 | - |
| SSU | CP075167.1 | 277,983 | 8,172 | 6,878 | - |
| SSU | CP075167.1 | 277,983 | 270,062 | 271,356 | + |
| LSU | CP075167.1 | 277,983 | 271,465 | 273,846 | + |
| telomere | CP075167.1 | 277,983 |  | at end | + |
| telomere | CP075168.1 | 270,484 | at start |  | - |
| LSU | CP075168.1 | 270,484 | 6,742 | 4,366 | - |
| SSU | CP075168.1 | 270,484 | 8,146 | 6,852 | - |
| SSU | CP075168.1 | 270,484 | 262,090 | 263,383 | + |
| LSU | CP075168.1 | 270,484 | 263,493 | 265,865 | + |
| telomere | CP075168.1 | 270,484 |  | at end | + |
| ***G. vavrai*** |  |  |  |  |  |
| SSU | scaffold_1 | 1,930,326 | 103,982 | 105,300 | + |
| LSU | scaffold_1 | 1,930,326 | 105,429 | 107,905 | + |
| LSU | scaffold_1 | 1,930,326 | 142,978 | 140,505 | - |
| SSU | scaffold_1 | 1,930,326 | 144,425 | 143,107 | - |
| SSU | scaffold_2 | 1,437,478 | 271,366 | 272,684 | + |
| LSU | scaffold_2 | 1,437,478 | 272,813 | 275,288 | + |
| LSU | scaffold_2 | 1,437,478 | 305,412 | 302,667 | - |
| SSU | scaffold_2 | 1,437,478 | 306,859 | 305,541 | - |
| SSU | scaffold_5 | 1,140,537 | 233,026 | 234,344 | + |
| LSU | scaffold_5 | 1,140,537 | 234,473 | 236,949 | + |
| LSU | scaffold_5 | 1,140,537 | 268,787 | 266,311 | - |
| SSU | scaffold_5 | 1,140,537 | 270,234 | 268,916 | - |
| telomere | scaffold_9 | 986,934 | at start |  | - |
| SSU | scaffold_10 | 917,715 | 297,190 | 298,508 | + |
| LSU | scaffold_10 | 917,715 | 298,637 | 301,115 | + |
| LSU | scaffold_10 | 917,715 | 333,269 | 330,793 | - |
| SSU | scaffold_10 | 917,715 | 334,716 | 333,398 | - |
| SSU | scaffold_10 | 917,715 | 532,905 | 534,223 | + |
| LSU | scaffold_10 | 917,715 | 534,352 | 536,826 | + |
| LSU | scaffold_10 | 917,715 | 568,545 | 566,035 | - |
| SSU | scaffold_10 | 917,715 | 569,992 | 568,674 | - |
| SSU | scaffold_13 | 693,172 | 334,454 | 335,772 | + |
| LSU | scaffold_13 | 693,172 | 335,901 | 338,376 | + |
| LSU | scaffold_13 | 693,172 | 416,324 | 413,848 | - |
| SSU | scaffold_13 | 693,172 | 417,793 | 416,453 | - |
| SSU | scaffold_14 | 564,345 | 83,279 | 84,597 | + |
| LSU | scaffold_14 | 564,345 | 84,726 | 87,200 | + |
| LSU | scaffold_14 | 564,345 | 105,121 | 102,647 | - |
| SSU | scaffold_14 | 564,345 | 106,568 | 105,250 | - |
| ***G. intestinalis*** |  |  |  |  |  |
| telomere | scaffold_1 | 682,557 | at start |  | - |
| SSU | scaffold_1 | 682,557 | 14,901 | 13,643 | - |
| LSU | scaffold_1 | 682,557 | 17,566 | 15,212 | - |
| LSU | scaffold_1 | 682,557 | 676,179 | 678,533 | + |
| SSU | scaffold_1 | 682,557 | 678,844 | 680,102 | + |
| telomere | scaffold_1 | 682,557 |  | at end | + |
| telomere | scaffold_2 | 662,782 | at start |  | - |
| SSU | scaffold_2 | 662,782 | 26,854 | 25,596 | - |
| LSU | scaffold_2 | 662,782 | 29,519 | 27,165 | - |
| telomere | scaffold_2 | 662,782 |  | at end | + |
| SSU | scaffold_3 | 512,068 | 89 | 1 | - |
| LSU | scaffold_3 | 512,068 | 2,754 | 400 | - |
| telomere | scaffold_4 | 496,367 | at start |  | - |
| LSU | scaffold_4 | 496,367 | 3,891 | 1,537 | - |
| LSU | scaffold_4 | 496,367 | 467,075 | 469,429 | + |
| SSU | scaffold_4 | 496,367 | 469,740 | 470,998 | + |
| telomere | scaffold_4 | 496,367 |  | at end | + |
| telomere | scaffold_5 | 439,321 | at start |  | - |
| SSU | scaffold_5 | 439,321 | 21,072 | 19,814 | - |
| LSU | scaffold_5 | 439,321 | 23,737 | 21,383 | - |
| LSU | scaffold_5 | 439,321 | 417,427 | 419,781 | + |
| SSU | scaffold_5 | 439,321 | 420,092 | 421,350 | + |
| telomere | scaffold_6 | 358,051 | at start |  | - |
| LSU | scaffold_6 | 358,051 | 355,541 | 356,848 | + |
| LSU | scaffold_6 | 358,051 | 356,869 | 357,109 | + |
| telomere | scaffold_6 | 358,051 |  | at end | + |
| telomere | scaffold_7 | 313,543 |  | at end | + |
| telomere | scaffold_8 | 302,620 | at start |  | - |
| SSU | scaffold_8 | 302,620 | 5,474 | 4,216 | - |
| LSU | scaffold_8 | 302,620 | 8,143 | 5,789 | - |
| telomere | scaffold_8 | 302,620 |  | at end | + |
| telomere | scaffold_9 | 301,496 | at start |  | - |
| SSU | scaffold_9 | 301,496 | 19,927 | 18,669 | - |
| LSU | scaffold_9 | 301,496 | 22,592 | 20,238 | - |
| SSU | scaffold_9 | 301,496 | 48,989 | 47,731 | - |
| LSU | scaffold_9 | 301,496 | 51,654 | 49,300 | - |
| SSU | scaffold_9 | 301,496 | 79,845 | 78,587 | - |
| LSU | scaffold_9 | 301,496 | 82,510 | 80,156 | - |
| telomere | scaffold_9 | 301,496 |  | at end | + |
| SSU | scaffold_10 | 289,315 | 14,606 | 13,348 | - |
| LSU | scaffold_10 | 289,315 | 17,271 | 14,917 | - |
| telomere | scaffold_10 | 289,315 |  | at end | + |
| telomere | scaffold_11 | 269,236 | at start |  | - |
| SSU | scaffold_11 | 269,236 | 21,926 | 20,668 | - |
| LSU | scaffold_11 | 269,236 | 24,593 | 22,239 | - |
| LSU | scaffold_11 | 269,236 | 254,836 | 257,190 | + |
| SSU | scaffold_11 | 269,236 | 257,501 | 258,759 | + |
| telomere | scaffold_11 | 269,236 |  | at end | + |
| LSU | scaffold_12 | 260,371 | 227,543 | 229,897 | + |
| SSU | scaffold_12 | 260,371 | 230,208 | 231,466 | + |
| telomere | scaffold_12 | 260,371 |  | at end | + |
| telomere | scaffold_13 | 238,023 | at start |  | - |
| telomere | scaffold_13 | 238,023 |  | at end | + |
| ***H. tvaerminnensis*** | |  |  |  |  |
| telomere | scaffold_1 | 2,681,470 | at start |  | - |
| telomere | scaffold_2 | 1,973,229 | at start |  | - |
| SSU | scaffold_2 | 1,973,229 | 582,225 | 583,527 | + |
| LSU | scaffold_2 | 1,973,229 | 583,631 | 586,118 | + |
| LSU | scaffold_2 | 1,973,229 | 628,198 | 625,710 | - |
| SSU | scaffold_2 | 1,973,229 | 629,605 | 628,303 | - |
| SSU | scaffold_2 | 1,973,229 | 1,202,433 | 1,203,735 | + |
| LSU | scaffold_2 | 1,973,229 | 1,203,840 | 1,206,328 | + |
| LSU | scaffold_2 | 1,973,229 | 1,236,773 | 1,234,006 | - |
| SSU | scaffold_2 | 1,973,229 | 1,238,180 | 1,236,878 | - |
| telomere | scaffold_2 | 1,973,229 |  | at end | + |
| telomere | scaffold_3 | 1,839,609 | at start |  | - |
| telomere | scaffold_3 | 1,839,609 |  | at end | + |
| SSU | scaffold_5 | 1,464,916 | 884,776 | 886,078 | + |
| LSU | scaffold_5 | 1,464,916 | 886,183 | 888,671 | + |
| LSU | scaffold_5 | 1,464,916 | 917,664 | 915,176 | - |
| SSU | scaffold_5 | 1,464,916 | 919,071 | 917,769 | - |
| telomere | scaffold_6 | 1,444,059 | at start |  | - |
| telomere | scaffold_6 | 1,444,059 |  | at end | + |
| telomere | scaffold_7 | 1,438,879 | at start |  | - |
| telomere | scaffold_8 | 1,177,745 | at start |  | - |
| telomere | scaffold_8 | 1,177,745 |  | at end | + |
| telomere | scaffold_9 | 1,176,229 | at start |  | - |
| telomere | scaffold_9 | 1,176,229 |  | at end | + |
| telomere | scaffold_10 | 1,005,214 | at start |  | - |
| telomere | scaffold_10 | 1,005,214 |  | at end | + |
| telomere | scaffold_11 | 1,001,064 |  | at end | + |
| telomere | scaffold_12 | 969,012 | at start |  | - |
| telomere | scaffold_12 | 969,012 |  | at end | + |
| SSU | scaffold_13 | 915,172 | 604,006 | 605,308 | + |
| LSU | scaffold_13 | 915,172 | 605,413 | 607,901 | + |
| LSU | scaffold_13 | 915,172 | 640,105 | 637,617 | - |
| SSU | scaffold_13 | 915,172 | 641,512 | 640,210 | - |
| telomere | scaffold_13 | 915,172 |  | at end | + |
| telomere | scaffold_14 | 824,917 | at start |  | - |
| telomere | scaffold_14 | 824,917 |  | at end | + |
| telomere | scaffold_15 | 779,687 | at start |  | - |
| telomere | scaffold_15 | 779,687 |  | at end | + |
| telomere | scaffold_16 | 723,150 | at start |  | - |
| SSU | scaffold_16 | 723,150 | 341,836 | 343,140 | + |
| LSU | scaffold_16 | 723,150 | 343,245 | 345,733 | + |
| LSU | scaffold_16 | 723,150 | 375,840 | 373,352 | - |
| SSU | scaffold_16 | 723,150 | 377,249 | 375,945 | - |
| telomere | scaffold_16 | 723,150 |  | at end | + |
| telomere | scaffold_17 | 676,572 | at start |  | - |
| telomere | scaffold_17 | 676,572 |  | at end | + |
| ***M. daphniae*** |  |  |  |  |  |
| 5S rRNA | scaffold_1 | 1,022,311 | 84,968 | 85,086 | + |
| 5S rRNA | scaffold_1 | 1,022,311 | 458,145 | 458,263 | + |
| 5S rRNA | scaffold_1 | 1,022,311 | 476,509 | 476,627 | + |
| telomere | scaffold_1 | 1,022,311 |  | at end | + |
| 5S rRNA | scaffold_3 | 875,858 | 549,646 | 549,528 | - |
| 5S rRNA | scaffold_4 | 864,371 | 193,363 | 193,245 | - |
| 5S rRNA | scaffold_4 | 864,371 | 440,735 | 440,617 | - |
| 5S rRNA | scaffold_10 | 180,591 | 21,687 | 21,805 | + |
| ***O. colligata*** |  |  |  |  |  |
| LSU | scaffold_1 | 410,041 | 45,174 | 42,618 | - |
| SSU | scaffold_1 | 410,041 | 374,995 | 376,346 | + |
| LSU | scaffold_1 | 410,041 | 376,587 | 379,213 | + |
| LSU | scaffold_2 | 404,995 | 22,755 | 20,129 | - |
| SSU | scaffold_2 | 404,995 | 24,380 | 22,996 | - |
| SSU | scaffold_2 | 404,995 | 375,181 | 376,567 | + |
| LSU | scaffold_2 | 404,995 | 376,808 | 379,434 | + |
| LSU | scaffold_3 | 392,470 | 41,285 | 38,659 | - |
| SSU | scaffold_3 | 392,470 | 42,877 | 41,526 | - |
| SSU | scaffold_3 | 392,470 | 387,626 | 388,977 | + |
| LSU | scaffold_3 | 392,470 | 389,218 | 391,844 | + |
| LSU | scaffold_4 | 380,045 | 64,371 | 61,745 | - |
| SSU | scaffold_4 | 380,045 | 65,963 | 64,612 | - |
| SSU | scaffold_4 | 380,045 | 367,353 | 368,704 | + |
| LSU | scaffold_4 | 380,045 | 368,945 | 371,571 | + |
| LSU | scaffold_5 | 374,328 | 33,429 | 30,873 | - |
| SSU | scaffold_5 | 374,328 | 324,218 | 325,602 | + |
| LSU | scaffold_5 | 374,328 | 325,843 | 328,469 | + |
| telomere | scaffold_5 | 374,328 |  | at end | + |
| LSU | scaffold_6 | 340,929 | 25,732 | 23,106 | - |
| SSU | scaffold_6 | 340,929 | 27,359 | 25,973 | - |
| SSU | scaffold_6 | 340,929 | 290,500 | 291,884 | + |
| LSU | scaffold_6 | 340,929 | 292,125 | 294,751 | + |
| telomere | scaffold_6 | 340,929 |  | at end | + |
| LSU | scaffold_7 | 332,792 | 7,677 | 5,051 | - |
| SSU | scaffold_7 | 332,792 | 9,302 | 7,918 | - |
| SSU | scaffold_7 | 332,792 | 283,773 | 285,159 | + |
| LSU | scaffold_7 | 332,792 | 285,400 | 288,026 | + |
| LSU | scaffold_8 | 275,127 | 36,497 | 33,871 | - |
| SSU | scaffold_8 | 275,127 | 38,122 | 36,738 | - |
| LSU | scaffold_9 | 270,892 | 7,588 | 4,962 | - |
| SSU | scaffold_9 | 270,892 | 9,213 | 7,829 | - |
| SSU | scaffold_9 | 270,892 | 257,958 | 259,342 | + |
| LSU | scaffold_9 | 270,892 | 259,583 | 262,209 | + |
| ***V. necatrix*** |  |  |  |  |  |
| telomere | CP142726.1 | 1,624,402 | at start |  | - |
| SSU | CP142726.1 | 1,624,402 | 481,028 | 482,274 | + |
| LSU | CP142726.1 | 1,624,402 | 482,389 | 484,989 | + |
| telomere | CP142726.1 | 1,624,402 |  | at end | + |
| telomere | CP142727.1 | 1,491,498 | at start |  | - |
| SSU | CP142727.1 | 1,491,498 | 852,365 | 853,611 | + |
| LSU | CP142727.1 | 1,491,498 | 853,726 | 856,247 | + |
| telomere | CP142727.1 | 1,491,498 |  | at end | + |
| telomere | CP142728.1 | 1,467,808 | at start |  | - |
| SSU | CP142728.1 | 1,467,808 | 833,476 | 834,722 | + |
| LSU | CP142728.1 | 1,467,808 | 834,837 | 837,358 | + |
| LSU | CP142728.1 | 1,467,808 | 969,466 | 966,865 | - |
| SSU | CP142728.1 | 1,467,808 | 970,827 | 969,581 | - |
| SSU | CP142728.1 | 1,467,808 | 1,031,111 | 1,032,357 | + |
| LSU | CP142728.1 | 1,467,808 | 1,032,472 | 1,034,994 | + |
| telomere | CP142729.1 | 1,360,009 | at start |  | - |
| LSU | CP142729.1 | 1,360,009 | 802,525 | 799,997 | - |
| SSU | CP142729.1 | 1,360,009 | 803,886 | 802,640 | - |
| LSU | CP142729.1 | 1,360,009 | 838,196 | 835,668 | - |
| SSU | CP142729.1 | 1,360,009 | 839,557 | 838,311 | - |
| telomere | CP142729.1 | 1,360,009 |  | at end | + |
| telomere | CP142730.1 | 1,297,884 | at start |  | - |
| LSU | CP142730.1 | 1,297,884 | 151,379 | 148,812 | - |
| SSU | CP142730.1 | 1,297,884 | 152,741 | 151,494 | - |
| LSU | CP142730.1 | 1,297,884 | 201,860 | 199,293 | - |
| SSU | CP142730.1 | 1,297,884 | 203,221 | 201,975 | - |
| telomere | CP142730.1 | 1,297,884 |  | at end | + |
| telomere | CP142731.1 | 1,281,651 | at start |  | - |
| SSU | CP142731.1 | 1,281,651 | 547,381 | 548,628 | + |
| LSU | CP142731.1 | 1,281,651 | 548,743 | 551,309 | + |
| SSU | CP142731.1 | 1,281,651 | 695,105 | 696,351 | + |
| LSU | CP142731.1 | 1,281,651 | 696,467 | 699,020 | + |
| telomere | CP142731.1 | 1,281,651 |  | at end | + |
| telomere | CP142732.1 | 1,234,891 | at start |  | - |
| SSU | CP142732.1 | 1,234,891 | 325,979 | 327,225 | + |
| LSU | CP142732.1 | 1,234,891 | 327,340 | 329,868 | + |
| telomere | CP142732.1 | 1,234,891 |  | at end | + |
| telomere | CP142733.1 | 1,094,101 |  | at end | + |
| telomere | CP142734.1 | 1,111,211 | at start |  | - |
| telomere | CP142734.1 | 1,111,211 |  | at end | + |
| telomere | CP142735.1 | 1,069,813 | at start |  | - |
| LSU | CP142735.1 | 1,069,813 | 160,673 | 158,341 | - |
| SSU | CP142735.1 | 1,069,813 | 162,034 | 160,788 | - |
| telomere | CP142735.1 | 1,069,813 |  | at end | + |
| telomere | CP142736.1 | 979,666 | at start |  | - |
| SSU | CP142736.1 | 979,666 | 608,539 | 609,785 | + |
| LSU | CP142736.1 | 979,666 | 609,900 | 612,234 | + |
| SSU | CP142736.1 | 979,666 | 662,389 | 663,635 | + |
| LSU | CP142736.1 | 979,666 | 663,751 | 666,328 | + |
| SSU | CP142736.1 | 979,666 | 707,548 | 708,794 | + |
| LSU | CP142736.1 | 979,666 | 708,910 | 711,480 | + |
| telomere | CP142737.1 | 992,507 | at start |  | - |
| SSU | CP142737.1 | 992,507 | 493,951 | 495,197 | + |
| LSU | CP142737.1 | 992,507 | 495,313 | 497,948 | + |
| LSU | CP142737.1 | 992,507 | 721,648 | 719,108 | - |
| SSU | CP142737.1 | 992,507 | 723,009 | 721,763 | - |
| telomere | CP142737.1 | 992,507 |  | at end | + |

**Table S3: Detailed methylation annotation of microsporidia.** Genomic features of each species are sorted from most overrepresented (= hypermethylated) to most underrepresented (= hypomethylated). Asterisks denote levels of statistical significance obtained from genome ontology. Specifically, **^*^** indicates p < 0.05, **^**^** indicates p < 0.01, and **^***^** indicates p < 0.001.

| **Genomic feature per species** | **Total length (bp)** | **Observed overlap (bp)** | **Expected  overlap (bp)** | **Log Ratio Enrichment** | **Log *P*-value: – = hypermethylated + = hypomethylated** | |
| --- | --- | --- | --- | --- | --- | --- |
| ***H. tvaerminnensis*** | |  |  |  |  |  |
| DNA repeat | 1,560,788 | 746 | 339 | 0.79 | -202.4 | **^***^** |
| LINE repeat | 1,582,664 | 423 | 344 | 0.21 | -11.48 | **^***^** |
| Unknown repeat | 5,143,073 | 1,219 | 1,118 | 0.09 | -8 | **^***^** |
| LTR | 57,422 | 26 | 12 | 0.77 | -7.53 | **^***^** |
| Satellite repeat | 16,080 | 7 | 3 | 0.85 | -2.73 |  |
| RC repeat | 16,403 | 5 | 3 | 0.51 | -1.25 |  |
| rRNA | 39,231 | 10 | 8 | 0.22 | -1.05 |  |
| Low complexity | 107,195 | 5 | 23 | -1.75 | 13.75 | **^***^** |
| mRNA | 5,010,107 | 925 | 1,089 | -0.16 | 19.55 | **^***^** |
| Simple repeat | 240,514 | 8 | 52 | -1.87 | 31.3 | **^***^** |
| ***G. intestinalis*** |  |  |  |  |  |  |
| Unknown repeat | 851,514 | 541 | 230 | 0.86 | -200.41 | **^***^** |
| LINE repeat | 1,275,131 | 627 | 345 | 0.6 | -136.83 | **^***^** |
| rRNA | 61,603 | 103 | 16 | 1.86 | -107.06 | **^***^** |
| LTR | 128,421 | 40 | 34 | 0.16 | -1.58 |  |
| Low complexity | 3,703 | 0 | 1 | 0 | 1 |  |
| Simple repeat | 29,428 | 0 | 7 | -1.95 | 7.99 | **^***^** |
| mRNA | 2,627,278 | 488 | 711 | -0.38 | 77.31 | **^***^** |
| ***G. vavrai*** |  |  |  |  |  |  |
| LTR | 1,100,597 | 705 | 416 | 0.52 | -91.96 | **^***^** |
| rRNA | 55,238 | 68 | 20 | 1.22 | -35.97 | **^***^** |
| DNA repeat | 1,713,959 | 811 | 648 | 0.21 | -21.86 | **^***^** |
| mRNA | 3,459,466 | 1,390 | 1,309 | 0.06 | -4.91 | **^**^** |
| Unknown repeat | 1,191,871 | 475 | 451 | 0.05 | -1.9 |  |
| LINE repeat | 153,485 | 52 | 58 | -0.11 | 1.46 |  |
| Low complexity | 139,751 | 17 | 52 | -1.24 | 21.08 | **^***^** |
| Simple repeat | 267,176 | 15 | 101 | -1.98 | 62.19 | **^***^** |
| ***B. daphniae*** |  |  |  |  |  |  |
| rRNA | 12,010 | 37 | 2 | 2.92 | -72.86 | **^***^** |
| Unknown repeat | 1,303,814 | 326 | 237 | 0.30 | -17.66 | **^***^** |
| mRNA | 2,987,996 | 596 | 544 | 0.09 | -5.39 | **^**^** |
| LTR | 1,089,800 | 213 | 198 | 0.07 | -1.91 |  |
| DNA repeat | 698,756 | 107 | 127 | -0.21 | 4.41 | **^*^** |
| Low complexity | 44,855 | 2 | 8 | -1.39 | 4.44 | **^*^** |
| LINE repeat | 313,660 | 39 | 57 | -0.38 | 5.06 | **^**^** |
| Simple repeat | 96,325 | 1 | 17 | -2.83 | 14.72 | **^***^** |
| ***E. intestinalis*** |  |  |  |  |  |  |
| rRNA | 83,137 | 1,422 | 93 | 2.73 | -2,936.43 | **^***^** |
| Unknown repeat | 84,912 | 310 | 95 | 1.18 | -163.04 | **^***^** |
| Low complexity | 2,581 | 0 | 2 | -0.69 | 2.90 |  |
| Simple repeat | 15,295 | 0 | 17 | -2.83 | 17.22 | **^***^** |
| mRNA | 2,111,255 | 1,026 | 2,368 | -0.84 | 1,476.72 | **^***^** |
